# Exploring the molecular makeup of support cells in insect camera eyes

**DOI:** 10.1101/2023.07.19.549729

**Authors:** Shubham Rathore, Aaron Stahl, Joshua B. Benoit, Elke K. Buschbeck

## Abstract

Animals generally have either compound eyes, which have evolved repeatedly in different invertebrates, or camera eyes, which have evolved many times across the animal kingdom. Both eye types include two important kinds of cells: photoreceptor cells, which can be excited by light, and non-neuronal support cells (SupCs), which provide essential support to photoreceptors. Despite many examples of convergence in eye evolution, similarities in the gross developmental plan and molecular signatures have been discovered, even between phylogenetically distant and functionally different eye types. For this reason, a shared evolutionary origin has been considered for photoreceptors. In contrast, only a handful of studies, primarily on the compound eyes of *Drosophila melanogaster*, have demonstrated molecular similarities in SupCs. *D. melanogaster* SupCs (Semper cells and primary pigment cells) are specialized eye glia that share several molecular similarities with certain vertebrate eye glia, including Müller glia. This led us to speculate whether there are conserved molecular signatures of SupCs, even in functionally different eyes such as the image-forming larval camera eyes of the sunburst diving beetle *Thermonectus marmoratus*. To investigate this possibility, we used an in-depth comparative whole-tissue transcriptomics approach. Specifically, we dissected the larval principal camera eyes into SupC- and retina-containing regions and generated the respective transcriptomes. Our analysis revealed several conserved features of SupCs including enrichment of genes that are important for glial function (e.g. gap junction proteins such as innexin 3), glycogen production (glycogenin), and energy metabolism (glutamine synthetase 1 and 2). To evaluate the extent of conservation, we compared our transcriptomes with those of fly (Semper cells) and vertebrate (Müller glia) eye glia as well as respective retinas. *T. marmoratus* SupCs were found to have distinct genetic overlap with both fly and vertebrate eye glia. These results provide molecular evidence for the deep conservation of SupCs in addition to photoreceptor cells, raising essential questions about the evolutionary origin of eye-specific glia in animals.

## Introduction

Among the diversity of animal eyes, there are near perfect examples of both convergent and divergent evolution, but the structure–function relationships of eye components have confounded evolutionary biologists for centuries. Eye structure has evolved independently multiple times within the animal kingdom, often adapting to the specific ecological needs of the bearer [1, 2]. The simplest light-detecting organ likely consisted of a single light-sensitive photoreceptor cell accompanied by a pigmented support cell (SupC). This primordial photodetector is expected to have provided the animal with directional information and light sensitivity [3]. The animal kingdom today features an astonishing variety of eyes, from simpler pigment-cup eyes to more complex eyes with well-developed optical lenses. The latter can be broadly divided into two categories: a) compound eyes, which comprise multiple tightly organized light-detecting units called ommatidia that have evolved in invertebrates, and b) camera eyes, which are single image-forming structures that have evolved many times in both vertebrates and invertebrates [2]. Despite apparent differences in evolutionary history, structure, and function, these eye types have distinct parallels. For example, both eye types have functionally similar cell types that can be broadly classified as photoreceptor cells, which can be excited by light, and non-neuronal SupCs, which provide essential support to photoreceptors. Parallels also exist at the molecular level in the form of specific gene regulatory networks (GRNs) that are generally conserved in eyes [4–8].

In regard to eyes, GRNs are best understood for early developmental processes that give rise to photoreceptor cells, the evolutionary origin, development, molecular framework, and function of which have been extensively studied. Comparative studies of mice and the fruit fly *Drosophila melanogaster* have identified functionally conserved molecular components such as the *pax6* family of genes (eyeless/ *Eye* and twin of eyeless/ *Toy*) and their targets *sine oculus/So*, *eyes absent*/*Eya*, and *dachshund*/*Dach* [9–11], which regulate early eye development in both invertebrates and vertebrates. Similarly, the proneural gene *atonal/Ath5* is required for the determination of the first retinal neuronal cell type in the eyes of both invertebrates (R8 photoreceptor) and vertebrates (retinal ganglion cell) [12]. These examples highlight a small subset of studies that have identified key components of conserved ancient GRNs related to photoreceptor development. More recently, SupCs have also been demonstrated to be vitally important for eyes, providing glia-like structural, metabolic, trophic, and functional support to photoreceptors [13][14]. Although less studied, some broad functional and molecular similarities, including components of GRNs, have been identified between the SupCs of species with distinct evolutionary origins. For example, recent studies have suggested that the vertebrate retinal pigmented epithelium is functionally analogous to the interommatidial pigment cells of *D. melanogaster* compound eyes. Both tissues physically interact with photoreceptor cells and provide essential support through neurotransmitter storage/recycling, lipid metabolism, ion homeostasis, energy support, and neuroprotection [15–17]. A second critical cell type in *D. melanogaster* compound eyes is the lens-secreting Semper cells, which also provide important support to adjoining photoreceptor cells [17]. Both cell types share many features with arthropod glia.

There are several types of arthropod glia, the functional specialization of which depends on their location in the nervous system. In *D. melanogaster*, based on the constitutive expression of the pan glial transcription factor *repo*, six distinct glial subfamilies are currently recognized: perineurial glia, subperineurial glia, cortex glia, astrocyte-like glia, ensheathing glia, and wrapping glia (for details of each type, see [18]). Additionally, specific glial subtypes that do not express *repo*, such as larval midline glia and peripheral sheath cells, have also been identified [18]. Thus, similar to vertebrate glia, arthropod glia are complex and their identification requires a combined understanding of support function and molecular expression. Arthropod glia exhibit several conserved functional parallels with vertebrate glia, including blood–brain barrier (BBB) formation, axon guidance, provision of metabolic/ionic support to neurons, neurotransmitter storage/recycling, structural support, and osmoregulation [19]. These functional parallels also extend to glial properties that were previously considered vertebrate specific. For example, a single study in *D. melanogaster* has shown that *cut*-positive wrapping glia envelop adjoining peripheral axons in a manner similar to the myelination of oligodendrocytes in vertebrates [20]. Functional parallels have also been drawn between the BBBs of vertebrates and *D. melanogaster*. In both cases, glia allow the transfer of specific support molecules that are required by the underlying neurons while limiting exposure to circulating fluids [21].

In vertebrate eyes, the Müller glia span the entire depth of the retina and have been suggested to act as light guides for photoreceptor cells by reducing light scatter [22]. These highly branched cells fine tune photoreceptor activity by a) neurotransmitter recycling (e.g., by expressing glutamine synthetase *glul*), b) potassium (K^+^) spatial buffering (e.g., by expressing inward rectifying K^+^ channel *kir4.1*), a process by which Müller glia regulate the excitability of retinal neurons [23], and c) osmoregulation through water transport mediated by aquaporins such as *aqp4* [24, 25]. Notably, in *D. melanogaster*, Semper cells show enriched expression of orthologous genes *gs2*, *kir4.1*, and *drip*, which are likely to perform analogous support functions [14]. Additionally, these cells are specifically marked by the expression of the transcription factor Cut, which is also expressed in other *D. melanogaster* glia [20, 26]. These similarities indicate the presence of SupC-specific GRNs, which could provide a new model for discovering conserved fundamental processes that regulate glia–neuron interactions. To explore this exciting avenue, it must be established whether glia-like functions are conserved in the SupCs of arthropod eyes other than *D. melanogaster* compound eyes. A particularly good model for this purpose is the high-functioning camera eyes of the larvae of holometabolous insects such as the sunburst diving beetle *Thermonectus marmoratus*.

*T. marmoratus* is a predacious insect (family: Dytiscidae). As a larva, *T. marmoratus* possesses 12 single-chambered camera-type eyes, 6 on each side [27], of which, 4 are enlarged with a cylindrical shape and are known as principal eyes (Fig. 1A). These eyes have been studied extensively in our laboratory in regard to development [28], anatomy [29], physiology [30, 31], and optics [32, 33]. We have shown that these eyes provide exceptional vision, which can be analyzed using optics, physiology, and behavior [31, 33, 34]. We have developed some molecular tools for genetically manipulating these larvae [35], which can be applied to understand specific molecular functions in these complex camera eyes. The larvae belong to the first extant species known to possess bifocal lenses, which likely assist in estimating prey distance [33]. These complex lenses are partly secreted by a subset of SupCs that are similar to those in *D. melanogaster* and are located in the distal tubular region of the principal eyes [32] (Fig. 1B). Aside from their role in lens secretion, the molecular make-up and function of SupCs in these camera eyes remain unknown. However, owing to the shared gross eye development plan of *T. marmoratus* camera eyes and *D*. *melanogaster* compound eyes [28] as well as the expected compound eye ancestry of the camera eyes of holometabolous larvae [36], these eyes are an exciting system for exploring the role of conserved GRNs. Furthermore, based on their organization in the principal eyes, the lens-secreting SupCs closer to the lens are expected to be similar to *D. melanogaster* pigment cells, whereas the SupCs closer to the underlying photoreceptors are expected to be similar to *D. melanogaster* Semper cells (Fig. 1C). Lastly, based on histological observations, these SupCs send out projections that enwrap the photoreceptor cells and are therefore well positioned to perform glia-typical functions. One strength of this study system is that the SupC-enriched tubular region is anatomically distinct from the retina, which allows the eyes to be physically divided into these two regions (Fig. 1D). This phenomenon provides a unique opportunity to conduct tissue-specific transcriptomics to understand the molecular makeup of SupCs without utilizing complex techniques such as FACS sorting.

**Fig. 1:**
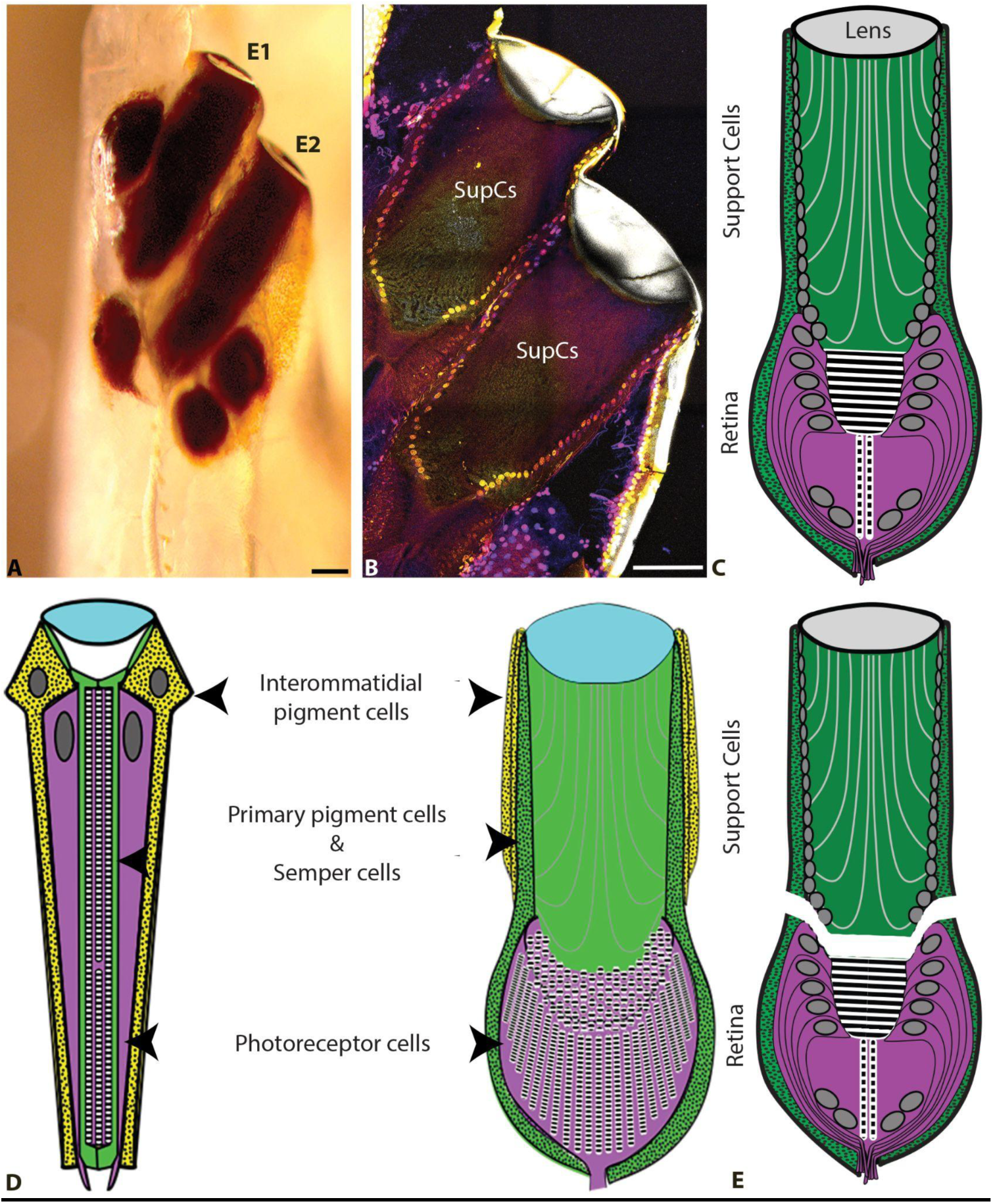
*T. marmoratus* larvae have two high-resolution image-forming principal camera eyes on each side of the head (E1 and E2). A. Both E1 and E2 are highly pigmented and tubular in shape, as illustrated by a freshly emerged larva in which the head cuticle is still transparent, scale bar = 100 µm. B. DAPI staining highlights the nuclei of the support cells (SupCs) that form the distal region of the eye tubes, scale bar = 100 µm. C. Schematic of a principal eye, illustrating its division into distally situated SupCs (green) and a proximal tiered retina (purple). D. These camera eyes and *D. melanogaster* compound eyes share similar developmental plans [28]. Based on their organization, it has been hypothesized that the outer SupCs in *T. marmoratus* camera eyes are related to *D. melanogaster* interommatidial pigment cells (yellow) and the inner SupCs to *D. melanogaster* primary pigment and Semper cells (green). E. As SupCs and photoreceptor cells are anatomically distinct, they can be dissected into separate regions for tissue-specific transcriptomics.

Here, we generated bulk SupC- and retina-specific transcriptomes to test the hypothesis that the SupCs in *T. marmoratus* camera eyes have glia-typical gene expression and function. It is important to point out that this method was unable to differentiate between specific SupC types; therefore, the resulting transcriptomes broadly reflect the SupC population. We also tested for overlap between the expression profiles of these SupCs and those of *D. melanogaster* Semper cells and zebrafish and mouse Müller glia to identify specific genes that could be part of generally conserved SupC-specific GRNs in animal eyes.

## Methods

### Animal husbandry, RNA isolation and RNA sequencing

All *T. marmoratus* larvae came from our lab-reared colony and were raised in a 14 h light–10 h dark cycle at 25°C. The larvae used in this study were 3–4 day old third instars. For RNA isolation, each individual was anesthetized on ice and dissected in RNA*later*^TM^ solution (Invitrogen, #AM7021). The two principal eyes (E1 and E2) pooled from 20–24 larvae were dissected into SupC-rich tubes and photoreceptor-rich regions, collected separately in RNA*later*^TM^ solution, and stored at -20°C until further processing. Three such biological replicates of the two tissues were generated. The total RNA from all tissues was isolated using an RNeasy Lipid Tissue Mini kit (Qiagen, #74804) according to the manufacturer’s protocol.

Quality control and sequencing of the eluted samples were performed by the DNA Sequencing and Genotyping Core at the Cincinnati Children’s Hospital Medical Center. Poly(A) libraries were prepared and sequenced on an Illumina NovaSeq 6000 system. For each sample, 20 million paired-end reads of 100 bp in length were generated. Raw reads for the support cell, retina and molting transcriptomes were deposited to the NCBI Sequence Read Archive, project numbers PRJNA995340 and PRJNA995342 .

### De novo assembly

The raw RNA seq reads for each eye-specific sample as well as the *T. marmoratus* transcriptomes previously published by our group [32] were trimmed using default settings and assessed for quality using FastQC [37].

All datasets were combined (∼250 million reads) assembled *de novo* with both CLC genomics workbench 12 (Qiagen, 12.0) and Trinity using default settings. The two assemblies were combined to generate the final contig library. Duplicate reads in the assembly were eliminated using CD-HIT [38, 39]. TransDE was used to determine open reading frames for protein coding genes and the assembly and the CDS version was annotated with the *D.melanogaster* proteome (Contigs with an e value less than 10^-10^ and a bit score over 80 were considered a match to Drosophila). Completeness of the transcriptome was estimated with BUSCO (version 4) [40], which was at 98.2% (complete)

### DE seq analysis

To identify the genes enriched in the SupC and retina regions, differential RNA seq analysis was performed on the CLC Genomics Workbench (Qiagen, 12.0) with the default settings, previously used in other arthropod systems [41–43]. The reads were normalized to transcripts per million (TPM). Statistical analyses were performed using an EDGE test and a Bbaggerly’s test (specifically for D.melanogaster due to single replicates) to identify the transcripts that were significantly enriched in the two tissues using a false discovery rate (FDR) adjusted p-value cut-off of <0.02. To identify key tissue-specific biological processes, GO analysis was performed on g:Profiler using the *D. melanogaster* as a proxy for the corresponding *T. marmoratus* contigs in each transcriptome [44]. Treemaps were constructed for both tissues based on enriched GO categories with Revigo[45].

### Transcriptome validation and heat map generation for glia-like genes

To validate the SupC transcriptomes, tissue-specific enriched genes were compared with the gene list of 10 *T. marmoratus* lens proteins previously identified as contributing to the lens [32], which itself has been shown to arise through secretion by the SupCs [28]. Contigs with a BLAST e-value less than 10^-10^ and an FDR-adjusted p-value of <0.02 were selected. Validation of the retina transcriptomes was performed using contig cut-offs similar to those described for the SupCs. Genes known to be related to photodetection and transduction were selected for validation. To investigate the possibility that the SupCs could be glial in nature, genes known from other glia or associated with glia-typical support functions were selected based on the above-mentioned parameters. Expressed genes that did not show significant enrichment in either tissue were also included. The normalized gene expression values for the selected genes were plotted as heat maps on R using R studio with the pheatmap package [46].

### Interspecies comparison

For interspecies comparisons, *D. melanogaster* Semper cell and photoreceptor transcriptomes [17] along with mouse and zebrafish Müller glia and retina neuron transcriptomes [47] were downloaded from the NCBI Sequence Read Archive (SRA). Raw files of these RNA-seq datasets were treated as described above to establish tissue-specific enrichment To assess overlap, each transcriptome was compared with the *T. marmoratus* transcriptomes using the BLASTx function. Overlapping *T. marmoratus* contigs were identified based on BLAST e-value cut-offs of 10^-100^ for *D. melanogaster* transcriptomes and 10^-60^ for mouse and zebrafish transcriptomes. We compared the tissue-specific enriched transcriptomes in all permutations and combinations between all tissue types (SupCs, Semper cells, Müller glia, retina, photoreceptors, and retinal neurons) using a freely available Venn diagram software (https://bioinformatics.psb.ugent.be/webtools/Venn/). Contigs that overlapped in the SupCs of all species and in the retinal cells of all species were annotated based on the *D. melanogaster* proteome, and the associated function in *D. melanogaster* was listed as indicated on FlyBase [48]

### Immunohistochemistry

Cut antibody staining was performed using a protocol modified from [8]. *T. marmoratus* third instar larvae were dissected and fixed in 4% formaldehyde solution, washed with PBS, and flash frozen in Neg50. The heads were cryosectioned sagittally into ∼20 µm slices on a cryostat (Leica CM1850) and stained with an anti-Cut antibody (1:50; DSHB) overnight at 4°C. These slices were then stained with a secondary antibody (anti-mouse Alexa Fluor 488, Thermo Fisher #A32723), mounted in Fluoromount with DAPI (Thermo Fisher #00495952), and imaged with a Leica SP8 confocal microscope.

## Results

### Anatomically distinct SupC and retina regions show functional specialization at the molecular level

To understand whether the SupC and retina regions of *T. marmoratus* principal eyes are functionally distinct, we generated transcriptomes (see STab1 for expression levels of genes described in this manuscript) and characterized the contigs with enriched transcript levels for both tissues using gene ontology (GO). Based on the significant GO classes obtained, the SupCs were enriched in genes associated with the development of anatomical structures, regulation of various biological processes, translation initiation, response to external stimuli, oxoacid catabolism, and molting (Fig. 2A). Other minor groups included genes associated with developmental processes, multicellular organismal process, localization, small molecule metabolism, carbohydrate metabolism, and general cellular processes (Fig. 2A). In contrast, the retina region was enriched in genes related to transport regulation, export from cells, response to light stimuli, cell signaling, cell junction organization, nervous system processes, and lipid metabolism (Fig. 2B). Other minor groups included genes associated with cell communication, cell localization, cell signaling, responses to stimuli, locomotion, homeostasis, cell regulation, rhodopsin metabolism, circadian rhythm, development, and organic hydroxy compound metabolism (Fig. 2C). Taken together, the expression profiles of these two tissues suggest functional specialization, with SupCs regulating development and support functions and the retina region being involved in light detection and neuron-typical regulatory functions.

**Fig. 2:**
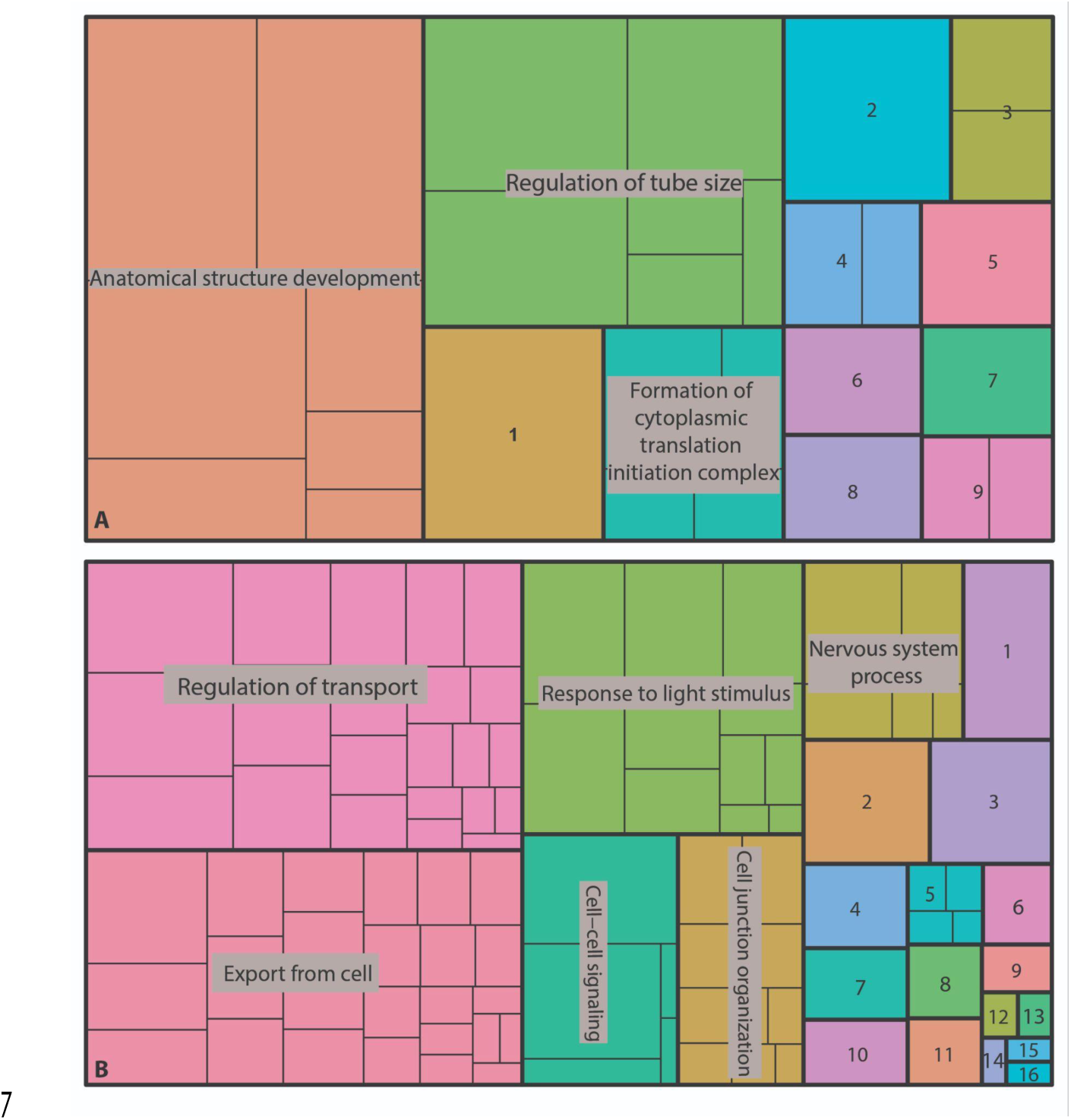
Validation of SupC- and retina-specific transcriptomes. A and B. Treemaps illustrating gene ontology (GO) terms for biological processes. A. SupCs are particularly enriched in genes from three major functional categories: anatomical structure development, tube size regulation, and cytosolic initiation complex formation. Additional minor categories include (1) cell developmental processes, (2) multicellular organismal processes, (3) molting, (4) response to external stimuli, (5) small molecule metabolic processes, (6) cellular localization, (7) carbohydrate metabolic processes, (8) cellular processes, and (9) oxoacid metabolic processes. B. The major functional categories in the retina are transport regulation, export from cells, response to light stimulus, neuron system processes, cell–cell signaling, and cell junction organization. Minor categories in this tissue include (1) cell communication, (2) cell localization, (3) cell signaling, (4) response to stimuli, (5) cellular lipid metabolic processes, (6) cell processes, (7) cell locomotion, (8)ell homeostatic processes, (9) rhodopsin metabolic processes, (10) multicellular organismal process, (11) biological regulation, (12) circadian rhythm, (13) rhythmic processes, (14) lipid metabolic process, (15) developmental processes and (16) organic hydroxy compound metabolic process.

### Validation of SupC and retina transcriptomes

To validate whether these two types of transcriptomes capture the expression of specific genes, we assessed the transcript levels of specific proteins that are expected to be enriched in each cell type. For SupCs, we evaluated the expression of lens protein genes, which has previously been characterized [32]. *T. marmoratus* larvae have structurally complex bifocal lenses that are partly secreted by the SupCs in the principal eyes. We previously identified 10 cuticular lens proteins that are enriched in the SupCs of the principal eyes, among which, only 2 (lens 6 and 7) also show some expression in the retina region [32]. Upon comparing the transcriptomes with the nucleotide sequences of the 10 lens proteins, we found 7 lens proteins that matched closely with respective contigs. Of these, lens 1, 2, 3, 5, 9, and 10 were exclusively enriched in the SupCs, whereas for lens 7, one of six contigs was enriched in the retina tissue (Fig. 3A). Other lens proteins (lens 4, 6, and 8) were not expressed highly enough to be detected in this analysis.

**Fig. 3:**
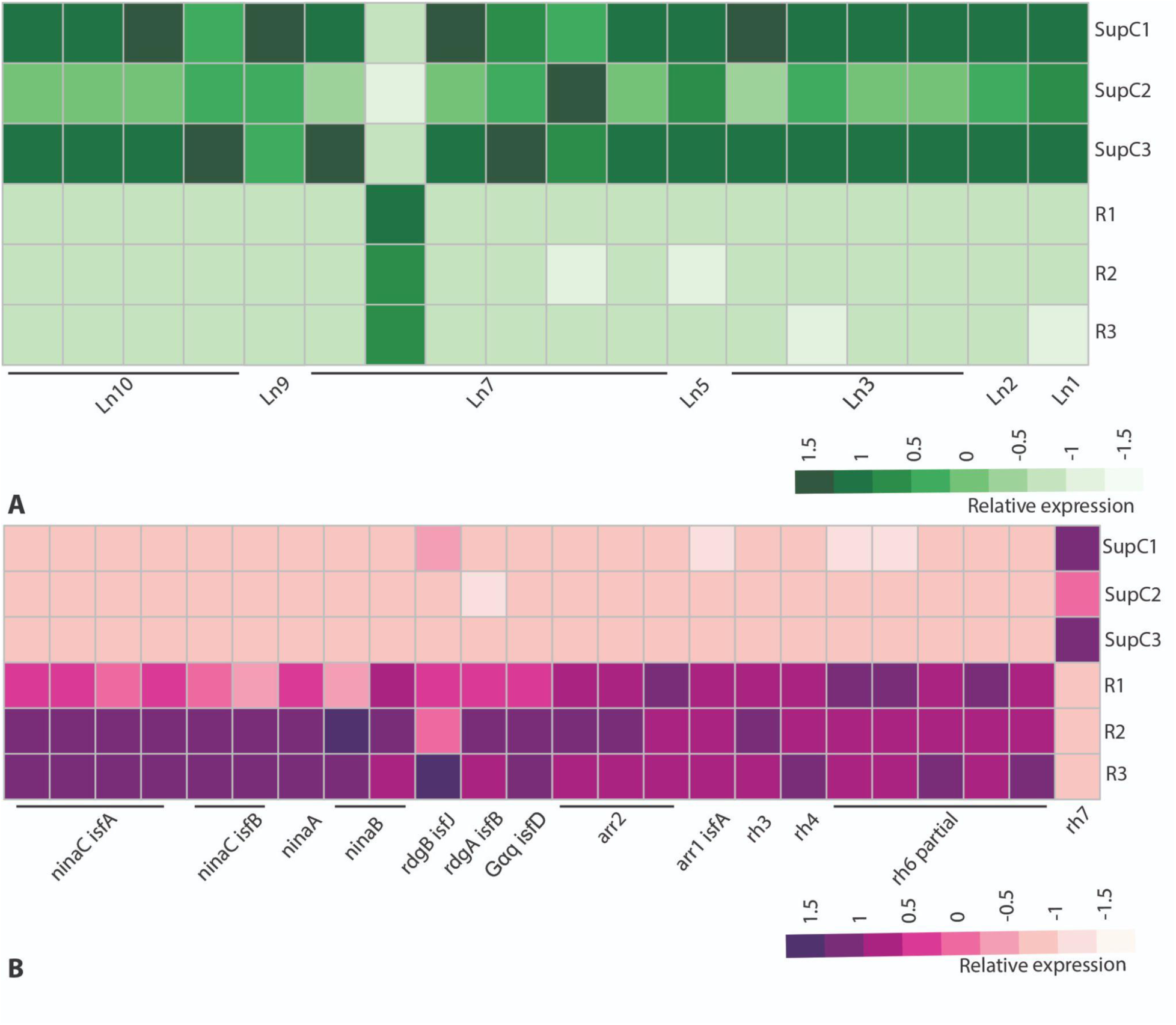
Heat maps represent gene expression, where darker colors indicate higher expression levels. A. As expected for this cell class, SupC transcriptomes are enriched in key lens (Ln) proteins (Ln 1–3, 5, 7, 9, and 10), with only one of six Ln 7 contigs being enriched in the retina transcriptomes. B. As expected, the retina transcriptomes are enriched in genes related to phototransduction and reception. These genes include arrestins 1 (*arr1* and 2 (*arr2*), retinal degeneration enzymes a (*rdgA*) and b (*rdgB*), neither inactivation nor afterpotential (*ninaA*, *ninaB*, and *ninaC*), and opsins *rh3*, *4*, and *6*. In contrast, nonvisual *rh7* is enriched in the SupCs.

For the retina, based on our previous work on photoreceptor cell development, anatomy, and opsin expression [27–29], we analyzed the expression of genes associated with photodetection and transduction. As the *T. marmoratus* contigs were annotated with the *D. melanogaster* proteome, the gene names used hereinafter follow the same nomenclature. We found three visual opsins, *rh4*, *rh3* (UV sensitive), and *rh6*-partial (green sensitive) [49], enriched in the retina, which agrees with a previous expression analysis [29]. In addition, a single nonvisual opsin, *rh7* [50], was enriched in the SupCs (Fig. 3B). Furthermore, typical invertebrate phototransduction genes such as *ninaC* isoforms a and b, *ninaA*, *ninaB*, *rdgA*, *rdgB*, *gqA*, and *arrestins* 1 and 2 [51] were enriched in the retina (Fig. 3B). Together, these results validated that the anatomical regions consisted predominantly of the expected cell types.

### Cut expression is conserved in a subset of SupCs

Several transcription factors showed either a tendency towards enrichment (cut *(ct),* bar homolog2 *(bh2)*) or significant enrichment (*eyes absent* (*eya*) and *sine oculis*) in the SupCs, whereas prospero (*pros*) showed a tendency towards enrichment in the retina (Fig. 4A). Of particular interest is the homeobox transcription factor Cut, which is expressed in many glial cell types in *D. melanogaster*, such as wrapping glia, sheath glia, and Semper cells [17, 18, 20], and is also conserved in the Semper cells of adult *T. marmoratus* compound eyes [8]. Therefore, Cut is a good candidate protein for confirming whether the larval *T. marmoratus* SupCs have deeply conserved molecular similarities with the Semper cells of *D. melanogaster* compound eyes (Fig. 1D), as previously proposed [28]. Specifically the findings are consistent with our homology model for insect ommatidia and *T. marmoratus* principal eyes. Conserved expression was only expected in a specific subset of SupCs, which in addition to generally low expression levels and a possible disjunction between the transcript and protein levels [52], could explain the relatively poor signal-to-noise ratio for *cut* expression. To investigate this possibility further, we used a *D. melanogaster* anti-Cut antibody, which was previously established to cross-react with *T. marmoratus* adult eyes [8] to stain cryosectioned larval camera eyes. Consistent with our predictions, Cut protein expression was sequestered in the nuclei of a small subset of distal SupCs in the principal eye tubes (Fig. 4B, arrowheads, and C, green nuclei). These results indicate that the small subset of Cut-positive SupCs are homologous to *D. melanogaster* Semper cells, thereby evidencing that at least this region of the eye tubes can serve as glia.

**Fig. 4:**
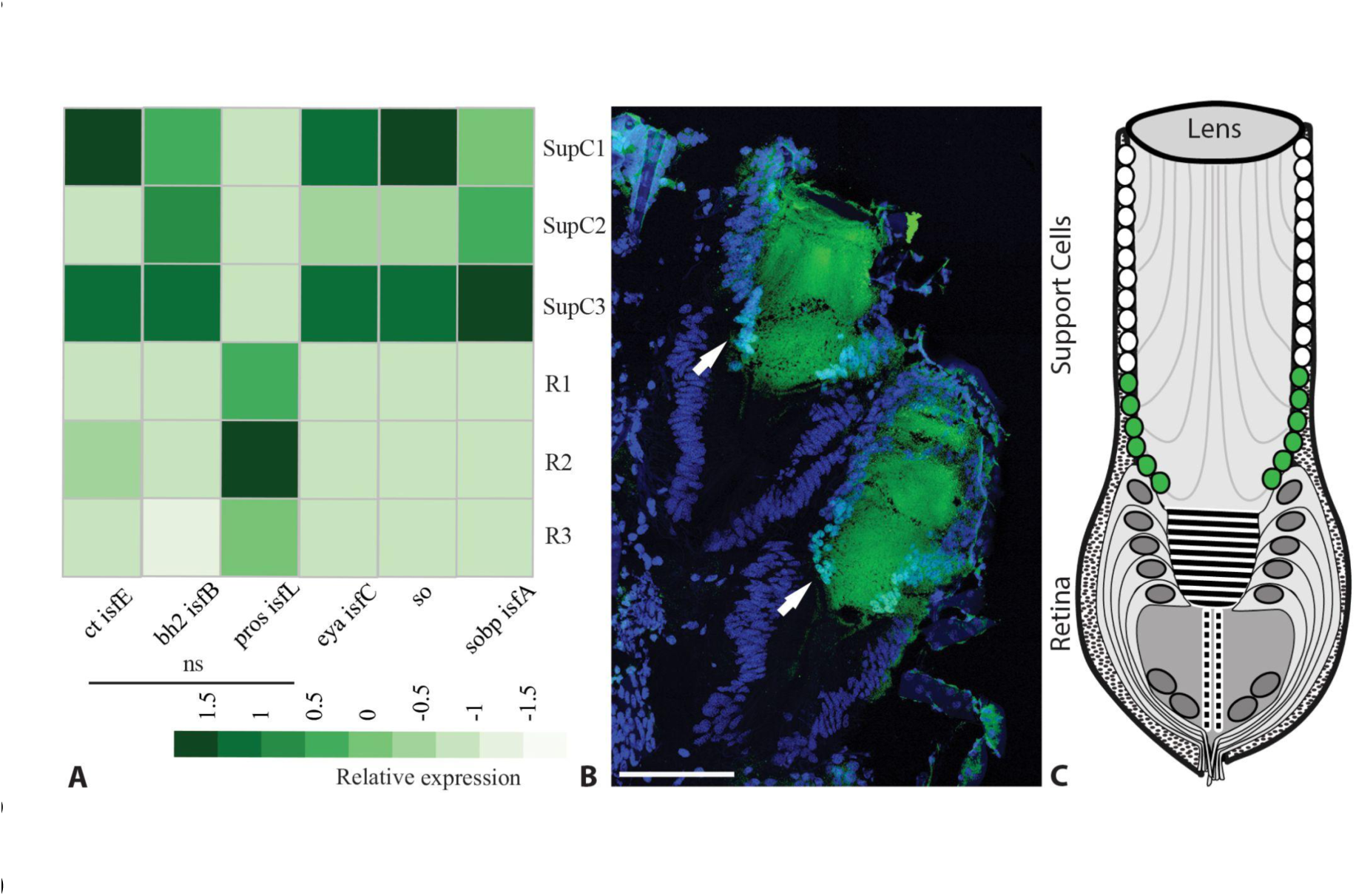
Expression of important transcription factors in the principal camera eyes of *T. marmoratus* third instars. A. Relative expression of transcription factors cut (*ct*) and bar homolog2 (*bh2*) shows a tendency towards but no significant enrichment in the SupCs. Conversely, prospero (*pros*) shows a tendency towards but no significant enrichment in the retina. In contrast, transcription factors *eyes absent* (*eya*) and *sine oculis* (so) as well as the sine oculis binding protein (*sobp*) are significantly enriched in the SupCs. B. Cut antibody, which is known to mark Semper cells in the compound eyes of *T. marmoratus* adults [8], stained a specific subset of proximally placed SupCs (teal, arrows), scale bar = 100 µm. C. As illustrated by the schematic, the staining pattern supports the deep conservation of this transcription factor and is consistent with our model, in which a portion of the SupCs in *T. marmoratus* larval eye tubes (green) are homologous to *D. melanogaster* Semper cells.

### Investigating glia-like support functions in the SupCs of camera eyes

Glia in both insects and vertebrates are known for their characteristic gene expression and associated support functions [18, 53]. To test whether SupCs could be glia, we assessed the SupC and retina transcriptomes for the enrichment of genes that are expressed in other insect glia. We found SupC enrichment of the following key glial genes (Fig. 5A): two isoforms (b and j) of myosin light chain kinase *strn-mlck*, which is expressed in *D. melanogaster* subperineurial glia and is essential for BBB integrity [54]; TGF-beta ligand *myo*, which is generally expressed in *D. melanogaster* glia and is necessary for neural circuit remodeling [55]; *axo*, a member of the neurexin superfamily that is expressed in ensheathing glia [56] and associated with neuronal excitability and synaptic plasticity [57]; and *ttk*, a C2H2 zinc finger domain transcription factor that is necessary for the differentiation of glia [18], including compound eye cone and Semper cells [58]. In contrast, *repo*, which is a marker for most glial cells in *D. melanogaster* [18], showed lower expression in the SupCs than in the retina region (Fig. 5A).

**Fig. 5:**
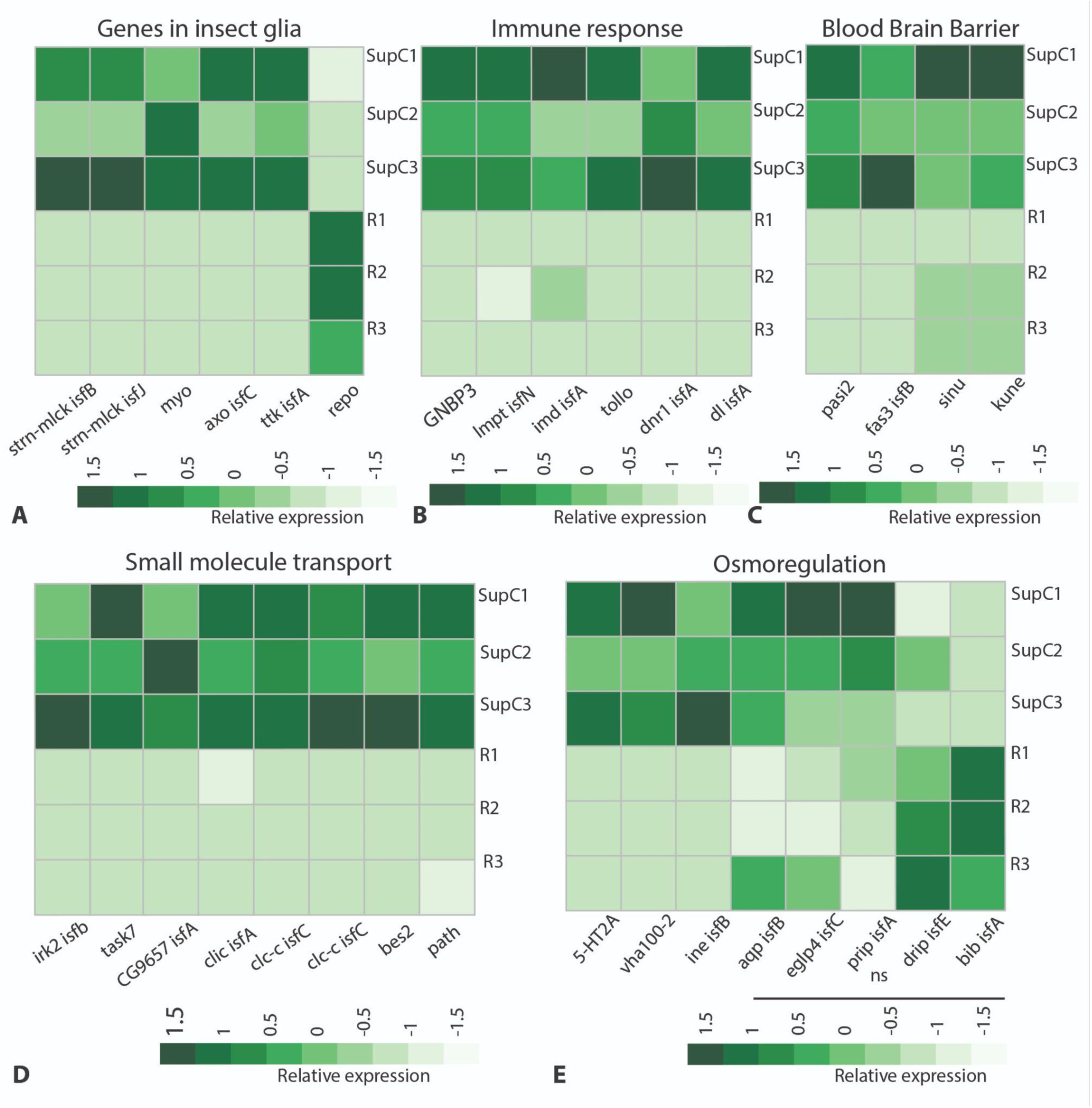
Homeostasis-related glia-typical functions in *T. marmoratus* camera eyes. A. Insect glia-typical genes such as stretchin-mick (*strn-Mlck*), myogialnin (*myo*), axotactin (*axo*), and tramtrack (*ttk*) are enriched in the SupCs, but the general insect glia marker reversed polarity (*repo*) is enriched in the retina. B. The SupCs are enriched in immune response genes, including gram-negative bacteria binding protein 3 (*gnbp3*), limpet (*lmpt*), immune deficiency (*imd*), Toll-like receptor (*tollo*), defense repressor 1 (*dnr1*), and dorsal (dl). C. SupC-enriched genes associated with blood–brain barrier (BBB) formation include pasiflora 2 (*pasi2*), fasciclin 3 isoform B (*fas3*), sinuous (*sinu*), and *kune*. D and E. Genes required for potassium transport (inwardly rectifying potassium channel 2 (*irk2*) and acid-sensitive potassium channel 7 (*task7*)), sodium symport (*rumpel/CG9657*), chloride transport (chloride channels (*clic* and *clc-c*) and bestrophin (*bes2*)), amine transport (pathetic (*path*)), osmoregulation (serotonin receptor (*5-ht2a*), vacuolar H+ ATPase (*vha100-2*), and osmotic stress response related gene inebriated (*ine*)) are enriched in the SupCs. The expression of aquaporin genes such as *aqp*, *eglp4*, *prip*, *drip*, and *bib* is not significantly different in the SupCs and retina.

Glia in *D. melanogaster* regulate innate immune responses to external antigens that are introduced as bacterial/fungal infections or traumatic injuries [59]. In the *T. marmoratus* principal eyes, the SupCs form a physical barrier between the external larval hemolymph and the internal photoreceptors, and this placement leads us to expect involvement in immune responses. Consistent with this expectation, we found SupC enrichment of bacterial and fungal response proteins such as *gnbp3*, *lmpt*, *imd*, and *tollo*. Additionally, ring domain ubiquitin ligase *dnr1* and transcription factor *dl*, which function downstream of Toll-like receptors (*tollo*), were also enriched in SupCs (Fig. 5B). Considering the vital role of *imd* and *tollo* in regulating the innate immune response in flies [59] and the expression of other immunity-related genes, the SupCs of *T. marmoratus* larval eye tubes are likely essential for regulating protective functions for the photoreceptors that they enwrap.

As in vertebrates, the central nervous system neurons of arthropods tend not to make direct contact with the hemolymph and are generally shielded by glia. Functions related to this BBB, including the presence of septate junctions between glial cells [60], are fundamental to certain glial cells and are best understood in *D. melanogaster*. We found SupC enrichment of the following septate junction formation genes: *pasi2*, *fas3*, *sinu*, and *kune* [61, 62] (Fig. 5C). Glia also mediate the transport of specific small molecules such as ions, other osmolytes, and water (Fig. 5D and E). Accordingly, we found SupC enrichment of small molecule transporters including potassium transporters (*irk2* and *task7*), a sodium symporter (rumpel/cg9657) that is also expressed in *D. melanogaster* ensheathing glia [63], chloride transporters (*clic*, *clc-c*, and *bes2*), and a glial amine transporter (*path*) [61] (Fig. 5D). For genes involved in ion transport and osmoregulation, we only found SupC enrichment of a serotonin receptor (*5-HT2A*) [64], a single vacuolar H+ ATPase (*vha100-2*), and a neurotransmitter transporter (*ine*) (Fig. 5E), which is expressed in the perineurial glia of *D. melanogaster* and is associated with water regulation in malpighian tubules (Luan et al., 2015). For other genes that are typically associated with water transport, including aquaporins *aqp*, *eglp4*, *prip*, *drip*, and *bib*, we did not find any significant enrichment in the SupCs compared with the retina region (Fig. 5E). In contrast, *drip* enrichment has been observed in *D. melanogaster* Semper cells [17]. Thus, there is transcriptomic support for SupCs playing a role in barrier formation between the neuronal part of the eye and its surroundings, including for ion transport, which may be associated with osmotic processes.

Glial cells also provide metabolic support to the adjoining neurons, which generally lack storage capabilities for energy-rich molecules such as carbohydrates and lipids [17, 65]. Hence, glucose uptake, transport, and storage are important support functions mediated by glia. Consistent with such functions, we found SupC enrichment of genes associated with the pentose phosphate pathway (PPP) (*zwi* and *pgd*) and glycogenesis (*Gbs76A*, *ABGE*, *gyg* isoforms I and B, and *atpcl*). In regard to glucose homeostasis, we found two glucose transporters (*pippin* and *glut1* isoform W) to be enriched in the SupCs, whereas two *glut1* isoforms (S and P) were enriched in the retina (Fig. 6A). Genes regulating glutamate metabolism such as *gs2* are consistently expressed in *D. melanogaster* astrocyte-like glia, ensheathing glia, and Semper cells [65]. Similarly, we found that two glutamate receptors were enriched in the SupCs (*Kair1D* and *clumsy*). Notably, a different glutamate receptor (*Ekar*) and a predicted glutamate receptor associated protein (*CG11155*) were enriched in the retina region (Fig. 6B). Lastly, the SupCs were enriched in both glutamine synthetase enzymes (*gs1* and *gs2*) (Fig. 6B), which further supports their glia-like nature.

**Fig. 6:**
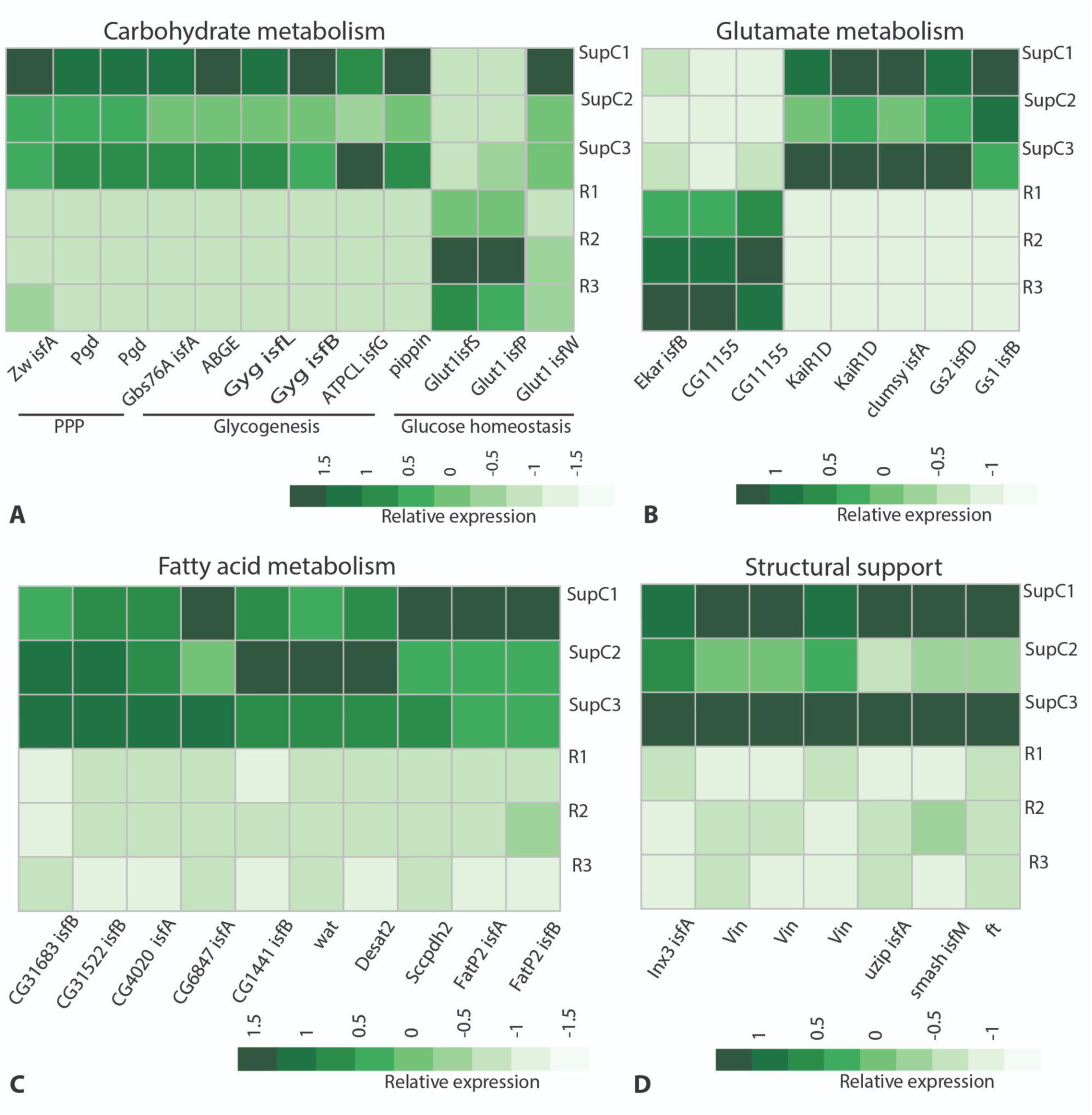
Metabolic and structurally related glia-typical support functions in *T. marmoratus* camera eyes. A. Genes associated with the pentose phosphate pathway (PPP), including zwischenferment (*zwi*), phosphogluconate dehyrogenase (*pgd*), and glycogen binding subunit 76A (*gbs76A*), and with glycogenesis, including 1,4-alpha-glucan branching enzyme (*abge*), glycogenin (*gyg*) isoforms I and B, and ATP citrate lyase (*atpcl*). For glucose homeostasis, we found glucose transmembrane transporter *pippin* and three isoforms of glucose transporter 1 (*glut1*); isoforms S and P are enriched in the retina region, whereas isoform W is enriched in the SupCs. B. A glutamate receptor and an associated protein (eye-enriched kainate receptor (*Ekar*)) as well as *CG11155* are enriched in the retina, whereas kainate-type ionotropic glutamate receptor subunit 1D (*Kair1D*) and a glutamate receptor activator *clumsy* are enriched in the SupCs. Glutamine synthetase enzymes *gs1* and *gs2* are also enriched in the SupCs. C. SupCs are enriched in several genes related to fatty acid metabolism including *CG31683*, *CG31522*, *CG4020*, *CG6847*, *CG1441*, waterproof (*wat*), desaturase 2 (*desat2*), saccherophin dehydrogenase 2 (*sccpdh2*), and fatty acid transporter protein 2 (*fatp2*). Structural support mediated by cell adhesion molecules is another important glial function. We found SupC enrichment of the following cell adhesion molecules: innexin 3 *(inx3*), vinculin (*vin*), unzipped (*uzip*), smallish (*smash*), and fat (*ft*).

Fatty acid storage and metabolism are another known support function of glial cells in the *D. melanogaster* nervous system [66]. Accordingly, we found SupC enrichment of many predicted lipid enzymes such as serine hydrolase *CG31683* isoform B, fatty acid elongase *CG31522* isoform B, fatty acyl co-A reductase *CG4020* isoform A, triacylglycerol lipase *CG6847* isoform A, and fatty acyl co-A reductase *CG1441* isoform B (Fig. 6C). Other lipid metabolism genes enriched in the SupCs included fatty acyl co-A reductase *wat*, which is necessary for tracheal lumen clearance [67], desaturase *desat2*, oxidoreductase *sccpdh2*, and two isoforms (A and B) of fatty acid transporter *fatP2* (Fig. 6C). Although these genes are associated with lipid metabolism, their role has not been tested directly in any *D. melanogaster* glia. Nevertheless, their consistent enrichment in the SupCs is suggestive of the nature of the metabolic pathways undertaken by these cells.

Finally, the SupCs were also enriched in cell adhesion molecules such as a gap junction protein (*inx3*), an alpha catenin family member (*vin*), a cell adhesion molecule expressed in longitudinal glia (*uzp*) [68], and an adherens junction associated protein (*smash*) (Fig. 6D). Cell adhesion is a key glial support function that is required by neurons for accurate structural development and function [69]. In addition, we also found an atypical cadherin family transmembrane protein (*ft*), which has a well-documented role in regulating hippo signaling and tissue growth [70]. However, it remains unclear whether *ft* plays a role in cell adhesion.

Overall, our expression analysis of the SupC region in the principal camera eyes of *T. marmoratus* provides clear evidence for the expression of many insect glia-typical genes and the enrichment of genes related to glia-typical structural, trophic, and metabolic support functions.

### Probing molecular overlap of functionally similar tissues in arthropod and vertebrate eyes

To determine whether any genes in the SupC and retina regions overlap with equivalent tissues in arthropod compound and vertebrate camera eyes, we adopted a comparative transcriptomics approach. For arthropods, we used the adult Semper cell and photoreceptor transcriptomes from [17], and for vertebrates, we used the control Müller glia and retinal neuron transcriptomes of mouse and zebrafish from [47]. There were 63 unique hits when contrasting SupCs and Semper cells, 20 unique hits when contrasting SupCs and Müller glia (zebrafish), and 31 unique hits when contrasting SupCs and Müller glia (mouse) with 6 genes that were common to all the species (Fig. 7A). These common genes are involved in triglyceride homeostasis, cuticular biosynthesis, glutamate metabolism, BMP signaling, and cell polarity maintenance (Fig. 7B). Similarly, there were 41 unique hits when contrasting retina and photoreceptors, 10 unique hits when contrasting retina and retinal neurons (zebrafish), and 37 unique hits when contrasting retina and retinal neurons (mouse) (Fig. 7C). An additional seven genes involved in glutamate response, enabling glutamate receptor activity, NMJ development, phototransduction, negative regulation of hippo signaling, and neurotransmitter secretion were common to all the species (Fig. 7D).

**Fig. 7:**
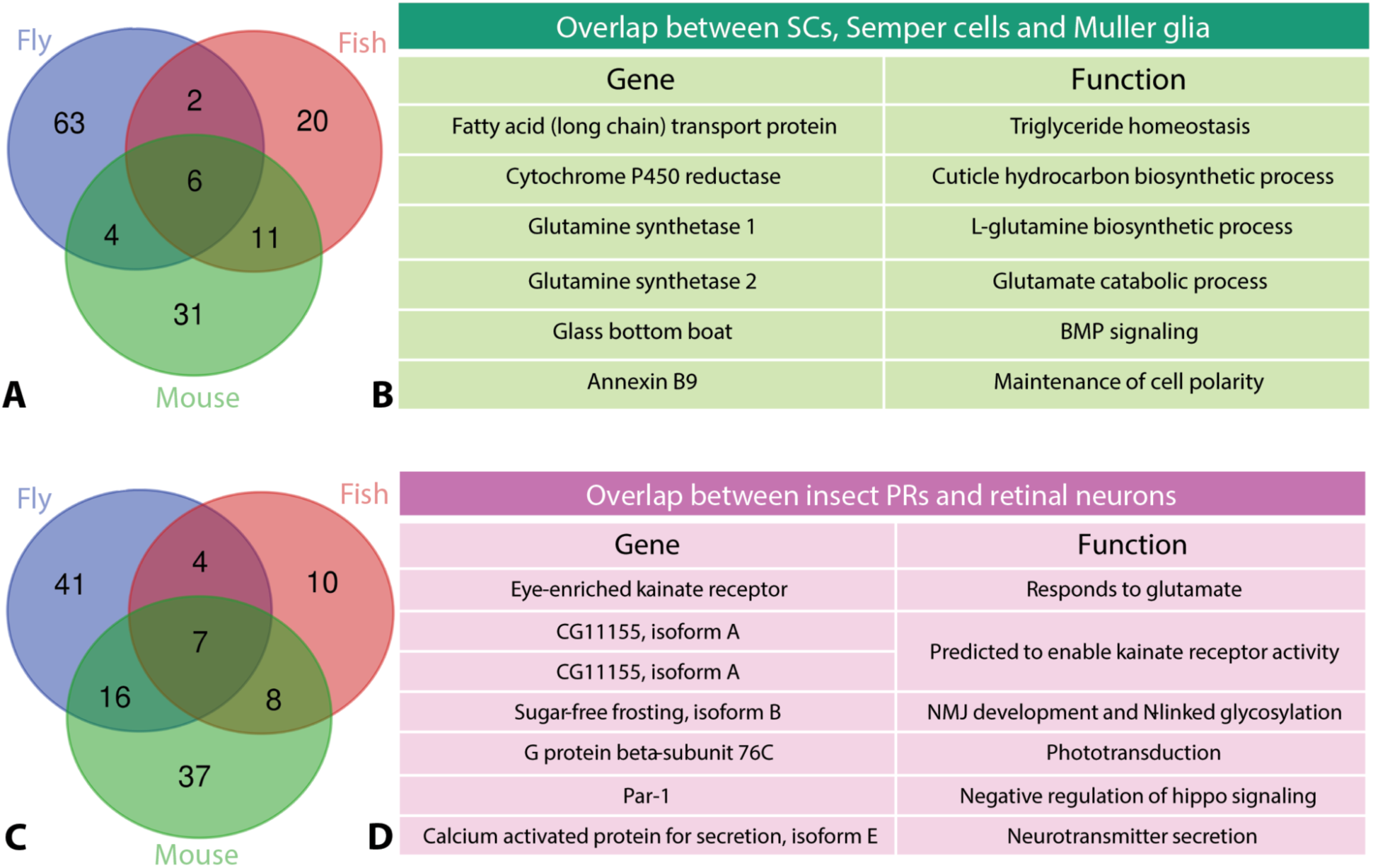
Four-way analysis to identify genes that overlap between the specific eye tissues of *T. marmoratus* and those of fly (*D. melanogaster*), fish (*D. rerio*), and mouse (*M. musculus*) [17, 47]. A. Comparison of *T. marmoratus* SupCs with *D. melanogaster* Semper cells and mouse and zebrafish Müller glia, revealing six common genes, the names and functions of which are listed in B. C. Comparison of the *T. marmoratus* retina with *D. melanogaster* photoreceptors cells and mouse and zebrafish retinal neurons, revealing seven common genes, the names and functions of which are listed in D.

## Discussion

Despite their distinct phylogenetic origins and functional specialization, different eye types have several conserved GRNs that are necessary for achieving important checkpoints during the formation of a functional eye [9–12]. The main objective of this study was to identify conserved GRNs in the relatively understudied SupCs of arthropod eyes, within the framework of camera eyes. The principal camera eyes of *T. marmoratus* larvae have well-developed SupCs that are anatomically distinct from the photoreceptors (Fig. 1B). Dissecting these eyes into proximal tubular SupC and distal retina regions presents a unique opportunity to explore SupC-specific molecular processes.

### Success of tissue-specific transcriptomics

The overall gene expression patterns are consistent with the successful separation of the larval camera eyes into SupC and retina regions. The molecular characterization clearly indicates tissue-specific functional specialization in these regions, with the SupC region regulating non-neuronal processes and the retina region showing enrichment in neuron-typical and photoreceptor-specific processes (Fig. 2). It should be noted that despite the clear functional distinction between the two regions, some level of cross-contamination between the tissues is expected. Specifically, as the SupCs have fine projections that wrap around the retina region, these cell portions were inadvertently included in the retina transcriptome. Conversely, a small medial retina in E1 extends into the SupC region [27]. Despite this limitation, our transcriptome verification is consistent with known functional specialization of the two eye regions. Consistent with our expectations, the SupCs are enriched in most of the lens proteins previously reported by our group (Stahl et al. 2017). Notably, for lens 7, one of the six contigs is enriched in the retina region (Fig. 3AB), in agreement with prior in situ staining, which suggests that lens 7 is expressed in both the SupC and photoreceptor regions [32]. Similarly, the enrichment of photodetection and transduction pathways in the retina region includes green-(*rh6*) and UV-sensitive visual opsins (*rh3* and *rh4*), which is consistent with the reported opsin expression [29] and spectral sensitivity [30] of *T. marmoratus* larvae (Fig. 3B). The only exception to the reported opsin expression in the SupCs is UV-sensitive nonvisual opsin *rh7*, which has not been previously described in *T. marmoratus*. However, Rh7 expression has been identified in the brain and compound eyes of *D. melanogaster* to generate a nonvisual photopigment [50, 71]. In the brain, Rh7 is expressed in a specific subset of neurons that are important for circadian rhythms [50]. In the compound eyes, some evidence suggests that Rh7 is involved in circadian entrainment [71], but otherwise its roles remain elusive. A reporter-based expression analysis revealed weak expression in R8 cells and high expression in the fenestrated layer, which consists of subretinal glia and pigment cells [72]. Although a deeper analysis of this expression pattern is needed in *D. melanogaster*, these data are consistent with our findings in the SupCs and retina of *T. marmoratus* camera eyes, suggesting the deep conservation of *rh7* in arthropods. Additionally, as the function of this opsin remains elusive in arthropod visual systems, it can be speculated that some of the SupC-specific processes related to circadian entrainment are regulated by *rh7*.

### Importance of Cut in a subset of SupCs

The expression of the transcription factor Cut in a subset of the SupCs in *T. marmoratus* (Fig. 4B and C) follows previous predictions regarding the cellular identity of specific regions in the complex principal eyes of *T. marmoratus* (Fig. 1D) and further suggests that the role of this transcription factor is conserved in arthropod eyes. We recently found that *cut RNAi* in the Semper cells of functionally different compound eyes (optical apposition eyes in *D. melanogaster* and optical superposition eyes in *T. marmoratus*) results in common deficits in the two eye types, including the general disorganization of the ommatidial array with incidences of lens fusion, lens defects, and rhabdomere displacement [8]. These observed parallels are insightful, as they point towards the conservation of key functions and provide an understanding of the role of Cut in functionally different contexts. The conservation of the role of Cut is also consistent with a general model for the development of most arthropod eyes [73], which includes predictions about how image-forming lens eyes may have evolved from compound eyes. These findings lead to questions regarding how a *cut* knockdown might affect the development and function of the *T. marmoratus* camera eyes. In fact, preliminary findings towards that end already suggest exciting parallels.

### Comparison of gene expression patterns in SupCs and retina generally support glial functions in SupCs

Identifying insect glia is a complex process due to their intricate molecular and functional profiles. Thus, it is expected that new glial subtypes are yet to be identified [18]. The transcription factor *repo* is a marker for most insect glial cells and is required for their differentiation but is not found in vertebrates [18]. Notably, not all glial cells in *D. melanogaster* are *repo*-positive [18], which allows insect glia to be divided into *repo*-expressing and non-expressing groups. For the Semper cells in *D. melanogaster*, *repo* expression is transient, only being detectable during the early developmental stages [17]. Therefore, it is plausible that the *T. marmoratus* SupCs do not show *repo* positivity because the transcriptomes are based on fully developed third instar larval eyes (Fig. 5A). The observed *repo* enrichment in the retina region may also be due to enrichment in the SupCs that tightly wrap around the photoreceptors. The enriched expression of insect glia genes other than *repo* in *T. marmoratus* SupCs (Fig. 5A) further supports our hypothesis that these cells are a type of glia.

The classification of *T. marmoratus* SupCs as glia is also evidenced by the gene expression patterns associated with specific support functions. The SupCs are enriched in genes associated with regulating homeostasis in the nervous system. Although only some of these genes are expressed in other insect glia, it could be speculated that the SupCs provide a broad range of homeostatic support to the adjoining photoreceptor cells, including the expression of genes that regulate immune responses in SupCs (Fig. 4B) and BBB-associated genes (Fig. 4C). Such functions could be important for SupCs because they directly interact with the hemolymph, which is the source of most pathogens. Similarly, the enriched expression of small molecule transport genes in the SupCs (Fig. 4D) suggests the active involvement of these cells in regulating neuronal access to specific molecules. Our data suggest that the SupCs also help maintain important ionic gradients around the photoreceptors. The enriched expression of inwardly rectifying potassium channel 2 (*irk2*) in the SupCs (Fig. 4D) is particularly notable, as this gene is also enriched in *D. melanogaster* Semper cells [17] and is homologous to vertebrate *Kir4.1*, which is required by Müller glia to maintain retinal function [24].

Another vital process that is relatively understudied in insect glia is osmoregulation. In vertebrate eyes, Müller glia provide osmoregulatory support to the photoreceptor cells by shuttling ions (with the help of ion channel *kir4.1*) and water (with the help of aquaporins such as *aqp4*) in and out of the eye [24]. In *D. melanogaster,* Semper cells also appear to be involved in these processes, as they are enriched in osmoregulatory genes including aquaporin *drip* [17], which shows sequence similarity with *aqp4*. However, *T. marmoratus* SupCs are only enriched in some related genes, such as those that support ion movements, with little enrichment in genes that code for water channels. Nevertheless, SupC enrichment of genes known to be important for osmoregulation in the malpighian tubules [64] is highly suggestive of an osmoregulatory role for these cells in the camera eyes of *T. marmoratus* larvae. Furthermore, the ubiquitous expression of aquaporins in the SupC and retina regions could highlight the importance of osmoregulation for the entire eye (Fig. 5E). Lastly, the lack of SupC-specific *aqp* enrichment could indicate that water channels are placed relatively permanently in these tissues, as is the case for malpighian tubules in *D. melanogaster* [74].

Glia also provide metabolic support to neurons [19]. Accordingly, the SupCs are enriched in genes that regulate carbohydrate, glutamate, and fatty acid metabolism (Fig. 6A–C), including the enrichment of glycogen storage genes. In contrast, glucose transporter *glut1* has a more complicated expression pattern, with isoform W being enriched in the SupCs and isoforms S and P being enriched in the retina region. These results are consistent with the known expression in both the glia [75], including Semper cells [17], and neurons [76] of *D. melanogaster*. The enhanced expression of PPP rate-limiting enzymes in the SupCs (Fig. 6A) is intriguing because this pathway is relatively understudied in *D. melanogaster* glia. A recent study found that the PPP is necessary for meeting the energetic needs of *D. melanogaster* neurons [75]. Additionally, the PPP is upregulated in vertebrate astrocytes in response to high glucose environments to combat oxidative stress [77]. Therefore, an understanding of the role of the PPP in arthropod glia could lead to important discoveries regarding the evolution of neuron–glia metabolic coupling. The enrichment of glutamine synthetases (*gs1* and *gs2*) in the SupCs and the expression of glutamate receptors in both the SupC and retina regions suggest a requirement for glutamate metabolism in these two tissues. Although this glial function is well understood in the glutamatergic synapses of both *D. melanogaster* [78] and vertebrates [79], it remains elusive in arthropod retina. Specifically, *gs2* is interesting because it is expressed in both *D. melanogaster* SupCs [17] and vertebrate Müller glia [80]. In Müller glia, this suggests metabolic rather than phototransduction processes, as *gs2* is required to maintain photoreceptor responses to light stimuli [81]. Similarly, as the arthropod retina uses histamine instead of glutamate as a neurotransmitter [82], it is likely that the glutamate– glutamine conversion reaction is also primarily related to metabolic support functions [14].

### Evolutionary implications

Despite considerable diversity in eyes, much discussion has focused on the deep conservation of photoreceptor cells [83], with relatively little attention given to other components. Our data suggest that certain functional roles of SupCs may also be deeply conserved. Specifically, our data point towards deeply conserved gene networks related to glia-typical functions within the SupCs of a high-functioning arthropod camera eye. In this context, *cut* is notable, as its expression is conserved in arthropod SupCs, even between functionally different compound and camera eyes. Further functional characterization will be helpful in advancing our understanding of SupC– photoreceptor coupling.

Some of the conserved functions are familiar from vertebrate eye glia and *D. melanogaster* Semper cells. Specifically, the six genes that overlap between *T. marmoratus* SupCs, *D. melanogaster* Semper cells, and mouse and zebrafish Müller glia (Fig.7 A & B) point towards ubiquitously important processes in phylogenetically and functionally different eye types. Notably, *gs2* is crucial because it is also conserved in vertebrate Müller glia and is necessary to maintain photoreceptor responses to light stimuli in rats [81]. Such findings can provide important insight into fundamental processes between eye-specific glia and adjacent photoreceptors. This is especially important from an evolutionary perspective, as the primordial photodetecting unit is thought of as a photoreceptor cell and a pigment cell [3]. These findings raise the possibility that such a simple visual organ might consist of an ancient, simplified neuron–glia pair. Deeply conserved SupC-specific genes could indicate the extent to which specific photoreceptor–SupC coupling processes are conserved. Finally, this work highlights how nontypical model organisms such as *T. marmoratus* larvae can be a powerful tool for discovering and understanding fundamental biological processes that are difficult to address in traditional model systems. Working with functionally different systems sheds light on how common molecular pathways could underlie the development of functionally different eyes. Expanding our comparisons beyond a handful of classical accessible model systems will allow us to capture more of the herculean biodiversity that is observed across arthropod visual systems and offer new insight into how visual systems have changed over evolutionary time.

## Supporting information

STable 1

## Acknowledgements

We would like to thank Dr. Tiffany A. Cook and Dr. Mark Charlton-Perkins for helpful guidance during the course of this project; the Genotyping and Sequencing Core at Cincinnati Children’s Hospital Medical Center for sequencing the transcriptomes; Chet Closson at the Live Microscopy Core (LMC, UCMC) for assistance with confocal imaging; Auggie Jester, Issac Wolff, Christine Swan, and Thiane Thiam for assistance with rearing the beetle larvae; and members of the Buschbeck laboratory for helpful suggestions.

## Funding sources

This research was funded by the National Science Foundation under IOS-1856241 (EKB), with partial support by NIH/NIAID grant R01AI148551 and NSF DEB-1654417 (JBB). A Weimann-Benedict Grant and Graduate Student Research Fellowship were awarded to SR by the Department of Biological Sciences and the Graduate Student Governance Association, University of Cincinnati.

## Author contributions

Conceptualization: EKB, SR

Data collection: SR, AS

Data curation: SR, JBB

Analysis of the transcriptome: SR, JBB

Analysis of Cut protein expression: SR, AS, EKB

Manuscript first draft: SR, EKB

Manuscript editing: SR, EKB, JBB, AS

## Notes

### Competing Interest Statement

The authors have declared no competing interest.

